# Oral prescription opioid-seeking behavior in male and female mice

**DOI:** 10.1101/621177

**Authors:** Alysabeth G. Phillips, Dillon J. McGovern, Soo Lee, Kyu Ro, David T. Huynh, Sophie K. Elvig, Katelynn N. Fegan, David H. Root

## Abstract

A significant portion of prescription opioid users self-administer orally rather than intravenously. Animal models of opioid addiction have demonstrated that intravenous cues are sufficient to cause drug-seeking. However, intravenous models may not model oral users, and the preference to self-administer orally appears to be partially influenced by the user’s sex. Our objectives were to determine whether oral opioid-associated cues are sufficient for relapse and whether sex differences exist in relapse susceptibility. Mice orally self-administered escalating doses of oxycodone under postprandial (prefed) or non-postprandial (no prefeeding) conditions. Both sexes demonstrated robust cue-induced reinstatement. In separate mice we found that oral oxycodone cues were sufficient to reinstate extinguished oral oxycodone-seeking behavior in the absence of postprandial or prior water self-administration training. During self-administration, we found that female mice earned significantly more mg/kg oxycodone than male mice. Follow-up studies indicated sex differences in psychomotor stimulation and plasma oxycodone/oxymorphone following oral oxycodone administration. In addition, gonadal steroid studies were performed in which we found divergent responses where ovariectomy enhanced and orchiectomy suppressed oral self-administration. While the suppressive effects of orchiectomy were identified across doses and postprandial conditions, the enhancing effects of ovariectomy were selective to non-postprandial conditions. These studies establish that 1) oral drug cues are sufficient to cause reinstatement that is independent of prandial conditions and water-seeking behavior, 2) earned oral oxycodone is larger in female mice compared with male mice potentially through differences in psychomotor stimulation and drug metabolism, and 3) gonadectomy produces divergent effects on oral oxycodone self-administration between sexes.

## Introduction

The United States is in the midst of an escalating prescription opioid epidemic (Buchanich, Balmert & Burke, 2017). Despite available opioid substitution treatments, individual differences exist with respect to which opioid addicted individuals will relapse (Bickel, Amass, Higgins et al., 1997; Epstein, Willner-Reid, Vahabzadeh et al., 2009; Preston, Kowalczyk, Phillips et al., 2017). Emerging evidence suggests that a primary aspect by which opioid addicted individuals differ is route of administration – oral versus intravenous (Back, Lawson, Singleton et al., 2011; Gasior, Bond & Malamut, 2016; Kirsh, Peppin & Coleman, 2012; Surratt, Kurtz & Cicero, 2011; Young, Havens & Leukefeld, 2010).

Those who orally abuse prescription opioids are significantly less likely to consume prescription or illicit opioids through intravenous, snorting, or smoking routes (Surratt, Kurtz & Cicero, 2011). While oral ingestion of prescription opioids has less immediate effects than intravenous use (Leow, Smith, Watt et al., 1992), oral oxycodone self-administration elicits dose-dependent subjective ratings of “liking” and “high” (Walsh, Nuzzo, Lofwall et al., 2008), and is a preferred route of administration (Gasior, Bond & Malamut, 2016; Kirsh, Peppin & Coleman, 2012; Surratt, Kurtz & Cicero, 2011). It has been reported that oral users circumvent the slower actions of oral opioids by chewing pills to increase the drug effects rate of onset (Gasior, Bond & Malamut, 2016; Kirsh, Peppin & Coleman, 2012). New abuse-deterrent reformulations of oxycodone have been designed to reduce chewing, snorting, or injecting prescription opioids but these formulations have resulted in an increase in oral oxycodone consumption (Cicero & Ellis, 2015; Gasior, Bond & Malamut, 2016).

Several individual differences influence oral prescription opioid use (Serdarevic, Striley & Cottler, 2017; Young, Havens & Leukefeld, 2010). Recent data suggests females are more likely to orally consume prescription opioids than use other routes of administration and women advance to regular prescription opioid use more quickly than males (Back, Lawson, Singleton et al., 2011; Gasior, Bond & Malamut, 2016; Kirsh, Peppin & Coleman, 2012). In rodent models, females show elevated oxycodone-induced locomotion (Collins, Reed, Zhang et al., 2016) and intravenous self-administration compared with males (Mavrikaki, Pravetoni, Page et al., 2017). While it is thought that sex hormones at least partially regulates drug-seeking behavior (Lynch, Roth & Carroll, 2002), it is unknown whether sex differences exist in oral oxycodone self-administration and if these potential differences are regulated by sex hormones. Together, oral consumption of prescription opioids is an important and increasing route of administration individuals use to self-administer. However, animal models of opioid addiction have largely focused on the intravenous route, which has limited our understanding of oral opioid use.

Environmental stimuli such as drug-associated cues influence relapse in opioid addicted individuals (Hyman, Fox, Hong et al., 2007; O’Brien, Childress, McLellan et al., 1990). In the clinic, intravenous users self-report significantly higher ratings of craving than oral users in response to drug-associated cues (McHugh, Fulciniti, Mashhoon et al., 2016; McHugh, Park & Weiss, 2014). These data predict that oral prescription opioid users are less susceptible to relapse triggered by drug-associated cues than intravenous users. Here, our objectives were to determine whether oral drug-associated cues are capable of causing relapse, and whether sex differences exist in the susceptibility to relapse, as modeled in the extinction-reinstatement paradigm (Shaham, Shalev, Lu et al., 2003). We also evaluated several factors that influence oral self-administration of prescription opioids between sexes.

## Materials and Methods

All animal procedures were performed in accordance with National Institutes of Health Guidelines and approved by the University of Colorado Institutional Animal Care and Use Committee.

### Cued reinstatement testing following oral self-administration

The self-administration paradigm was altered from a recently published procedure (Enga, Jackson, Damaj et al., 2016). Mice (8 male and 8 female C57BL6) were food restricted to 85% free-feeding body weight prior to and for the entirety of the experiment. Unless specified as non-postprandial, sessions consisted of mice receiving individual access to their daily food ration 1 hour prior to self-administration (postprandial conditions). Food remaining after the hour of isolated feeding was transferred to the self-administration chamber. If the ration was still not completely eaten after self-administration, it was returned with the mouse to the homecage following the session. Under non-postprandial conditions, mice were individually separated without their food ration 1 hour prior to self-administration and fed their daily ration in the home cage following self-administration. Mice self-administered water or oxycodone under postprandial or non-postprandial conditions by nose-poking an active hole (Table 1). All rewards were 20 μL. Inactive nosepokes were counted but had no programmed consequences. When mice satisfied the active nose-poke requirement for reward, the 7kHz tone cue was presented for 10 sec together with the termination of a light within the active hole.

**Table 1.** Experimental timeline. Mice self-administered water followed by escalating doses of post-prandial oxycodone and non-postprandial oxycodone. Following one week of abstinence, mice were tested for cue-induced reinstatement.

| Event | Oxycodone dose | Days | Postprandial |
| --- | --- | --- | --- |
| Water self-administration | 0.00 mg/ml | 3 | Yes |
| Oxycodone self-administration | 0.05 mg/ml | 3 | Yes |
| Oxycodone self-administration | 0.10 mg/ml | 3 | Yes |
| Oxycodone self-administration | 0.30 mg/ml | 3 | Yes |
| Oxycodone self-administration | 0.50 mg/ml | 3 | Yes |
| Oxycodone self-administration | 1.00 mg/ml | 3 | Yes |
| Oxycodone self-administration | 1.00 mg/ml | 7 | No |
| Abstinence | No oxycodone | 7 | No |
| Cue-induced reinstatement test | No oxycodone | 1 | No |

Mice initially self-administered water on a FR1 schedule of reinforcement until at least 10 active nosepokes were made in a single session. After water training mice self-administered at FR2. Mice self-administered water, 0.05 mg/ml oxycodone, 0.10 mg/ml oxycodone, 0.30 mg/ml oxycodone, 0.50 mg/ml oxycodone, and 1.00 mg/ml oxycodone under postprandial conditions for three days each (Table 1). Mice then self-administered 1.00 mg/ml oxycodone under non-postprandial conditions for seven days. After one week of forced abstinence mice were returned to self-administration chambers for reinstatement testing.

A between-within cue-induced reinstatement session (Shaham, Shalev, Lu et al., 2003) was used, consisting of an extinction phase and a cue phase (Root, Fabbricatore, Barker et al., 2009). The session began with the extinction phase. During the extinction phase, active and inactive nosepokes were counted but had no other programmed consequences. When mice emitted three or fewer active nosepokes over thirty consecutive minutes, the extinction criterion was met. When the extinction criterion was met the extinction phase ended and the cue phase began. During the cue phase, single active nosepokes activated the cue previously associated with oxycodone reward consisting of a ten second 7 kHz tone together with the termination of a light within the active hole. Inactive nosepokes were counted without other programmed consequence. Mice remained in the cue phase for 1 h.

### Cued reinstatement without postprandial conditions and prior water training

Mice (3 male and 3 female C57BL6) were food restricted to 85% free-feeding body weight prior to and for the entirety of the experiment. Mice non-postprandially self-administered 0.05 mg/ml on a FR1 schedule of reinforcement until at least 10 active nosepokes were made. Thereafter mice self-administered at FR2. Self-administration commenced non-postprandially for 0.05 mg/ml and 0.10 mg/ml over three days at each dose. After one week of forced abstinence mice were returned to self-administration chambers for the between-within cue-induced reinstatement session as described above.

### Psychomotor stimulation

Mice (6 male and 5 female C57BL6) were food restricted to 85% free-feeding body weight prior to and for the entirety of the experiment. Mice were orally administered 200 μL oxycodone at 8 mg/ml non-postprandially, three days later administered 16 mg/ml non-postprandially, three days later administered 16 mg/ml postprandially, and six days later administered 16 mg/ml non-postprandially again (Table 2). Doses were guided by prior research (Jacob, Poklis, Akbarali et al., 2017). Volume was chosen to mimic ten oral self-administration rewards. Immediately following administration, mice were placed in an open field (Stoelting AnyBox) for 20 min. Mice were videotracked by AnyMaze software (Stoelting) for 10 min, starting 3 min after placement into the open field, and average speed was calculated. Twenty minutes following injection, blood was collected.

**Table 2.** Experimental timeline. Mice were orally gavaged with different doses of oxycodone under non-postprandial (NPP) and postprandial (PP) conditions. Blood from the facial vein was collected twenty minutes following oxycodone administration.

| Event | Oxycodone dose | Days Between | Postprandial |
| --- | --- | --- | --- |
| Saline | 0.00 mg/ml | 3 | No |
| Oxycodone | 8 mg/ml | 3 | No |
| Oxycodone | 16 mg/ml | 3 | No |
| Oxycodone | 16 mg/ml | 3 | Yes |
| Oxycodone | 16 mg/ml | 6 | No |

### Detection of plasma oxycodone/oxymorphone

Twenty minutes after each oral oxycodone administration, whole blood was collected from the junction of the mandibular and facial veins (GoldenRod Lancet, Medipoint). Twenty minutes was chosen because brain concentrations plateau between 20 and 30 minutes following oral oxycodone administration (Jacob, Poklis, Akbarali et al., 2017). Approximately 100-500 μL whole blood was collected into ependorph tubes and subsequently centrifuged for 20 min at room temperature. The supernatant plasma was collected, frozen in liquid nitrogen, and stored at −80°C. Oxycodone/oxymorphone was detected in triplicate using ELISA (#221B-0096, Immunalysis) following the manufacturer’s instructions. Plasma samples, together with a standard curve of oxycodone in eight divisions between 0 and 20 ng/ml were read in a BioTek Elx808 microplate reader. Sample concentrations were calculated from an equation derived using a nonlinear least squares power fit of the standard curve (Matlab, Mathworks). Pilot experiments showed no detection of plasma oxycodone/oxymorphone from mice that did not receive oral oxycodone.

### Oral self-administration following gonadectomy

Mice (14 male and 18 female C57BL6) received either ovariectomy (n=9), orchiectomy (n=7), or sham (female=9, male=7) surgeries. After at least seven days of recovery, mice were food restricted to 85% free-feeding body weight for the entirety of the experiment. Mice then self-administered water and escalating doses of oxycodone as described in the *cued reinstatement testing following oral self*-*administration* section with the exceptions that mice self-administered non-postprandially for three days instead of seven and subsequently self-administered on a progressive ratio 4 schedule of reinforcement for three days non-postprandially (Table 3).

**Table 3.** Experimental timeline. Mice self-administered water followed by escalating doses of post-prandial oxycodone and non-postprandial oxycodone.

| Event | Oxycodone dose | Days | Postprandial |
| --- | --- | --- | --- |
| Water self-administration | 0.00 mg/ml | 3 | Yes |
| Oxycodone self-administration | 0.05 mg/ml | 3 | Yes |
| Oxycodone self-administration | 0.10 mg/ml | 3 | Yes |
| Oxycodone self-administration | 0.30 mg/ml | 3 | Yes |
| Oxycodone self-administration | 0.50 mg/ml | 3 | Yes |
| Oxycodone self-administration | 1.00 mg/ml | 3 | Yes |
| Oxycodone self-administration | 1.00 mg/ml | 3 | No |
| Progressive Ratio 4 | 1.00 mg/ml | 3 | No |

### Self-administration analyses

For analyses involving self-administration, including progressive ratio 4 performance, active nosepokes, inactive nosepokes, number of earned rewards, and earned mg/kg were averaged over the three days of self-administration at each dose and condition for each mouse. Earned mg/kg was defined as the number of rewards multiplied by the oxycodone mg dose in 20 μL volume, divided by kg body weight. Seven (dose) x two (sex) mixed ANOVAs were used to examine changes in active nosepokes, inactive nosepokes, rewards, and earned mg/kg per dose and sex (SPSS, IBM). If the assumption of sphericity was not met (Mauchley’s test), the Greenhouse–Geisser correction was used. Sidak-adjusted pairwise comparisons followed up significant main effects or interaction. Breakpoints were compared using independent t-tests for Gaussian distributed data or Mann Whitney tests in non-normally distributed data. In the experiment analyzing entirely non-postprandial self-administration without prior water training, data were non-normally distributed. Therefore, data comparing between two doses were analyzed using Wilcoxon signed ranks test (SPSS, IBM).

In the orchiectomy experiment, one sham mouse died overnight prior to non-postprandial testing. To statistically compare orchiectomy and sham male mice with a missing datapoint we used a mixed model repeated measures ANOVA with the non-postprandial datapoint of this single mouse defined as a missing data point (SPSS, IBM). The mixed model was also used to compare male and female sham mice. Covariance type was the diagonal matrix, the default covariance structure for repeated effects. Dependent and independent variables as well as posthoc Sidak-adjusted pairwise comparisons were identical. In mixed models, the denominator degrees of freedom are not integers but obtained by a Satterthwaite approximation. For the tests that used the mixed model reported here, denominator degrees of freedom were rounded to the closest integer.

### Cued reinstatement analyses

Active nosepokes/min and inactive nosepokes/min were calculated for the last thirty minutes of the extinction phase as well as the sixty minutes of the cue phase. Minute that mice reached extinction criterion was also determined. Extinction criterion time was analyzed by independent t-test between sexes. Two (phase) x two (sex) mixed ANOVAs were used to examine changes in active nosepokes/min and inactive nosepokes/min per phase and sex (SPSS, IBM). If the assumption of sphericity was not met (Mauchley’s test), the Greenhouse–Geisser correction was used. Sidak-adjusted pairwise comparisons followed up significant main effects or interaction. In the experiment analyzing entirely non-postprandial self-administration without prior water training, two reinstatement phases were compared using Wilcoxon signed ranks test (SPSS, IBM).

### Psychomotor stimulation and plasma oxycodone/oxymorphone analyses

Speed (mean meter/sec) was analyzed in two ways. We first used a mixed three (dose) x two (sex) ANOVA to examine changes in speed per dose and sex. Second, we used a mixed three (prandial condition) x two (sex) ANOVA to examine changes in speed per prandial condition and sex. Plasma oxycodone/oxymorphone was also analyzed for dose and prandial condition in two ways. First, a mixed two (dose) x two (sex) ANOVA was used to examine changes in plasma oxycodone/oxymorphone between 8 and 16 mg/ml doses and sex. Second, a mixed three (prandial condition) x two (sex) ANOVA was used to examine changes in plasma oxycodone/oxymorphone between prandial conditions at 16 mg/ml doses per sex. For all analyses, if the assumption of sphericity was not met (Mauchley’s test), the Greenhouse–Geisser correction was used. Sidak-adjusted pairwise comparisons followed up significant main effects or interaction.

## Results

Mice self-administered according to a modified version of a recently established oral oxycodone self-administration paradigm (Enga, Jackson, Damaj et al., 2016) (Table 1). Briefly, food restricted mice were fed their daily food ration prior to daily self-administration sessions for water or escalating doses of oxycodone (postprandial conditions). Mice then self-administered for one week under non-postprandial conditions at the highest oxycodone dose. Earned drug coincided with the sounding of a 10 sec tone and the termination of a light cue within the active hole.

Male and female mice reliably orally self-administered oxycodone across doses (Figure 1). Signs of intoxication, such as hyperlocomotion and Straub tail, were observed in all subjects, with higher doses most reliably eliciting these opioid-like behaviors. Mixed ANOVAs yielded significant main effects of dose on active nosepokes, F(6, 84) = 156.27, p < 0.001, inactive nosepokes, F(6, 84) = 3.49, p = 0.022, and rewards earned, F(6, 84) = 22.63, p < 0.001 without significant main effect of sex or dose X sex interaction. Posthoc Sidak-adjusted pairwise comparisons showed that when mice self-administered 1.00 mg/ml oxycodone under non-postprandial conditions, both active nosepokes and rewards earned decreased from 1.00 mg/ml oxycodone under postprandial conditions (p < 0.001) but inactive nosepokes were unchanged (p > 0.05) (Figure 1A-B). Examining earned drug normalized to the body weights of male and female mice (mg/kg) yielded a significant dose X sex interaction, F(6, 84) = 6.19, p < 0.01. Posthoc Sidak-adjusted pairwise comparisons showed that female mice had significantly higher earned mg/kg oxycodone between 0.30 and 1.00 mg/ml doses under postprandial conditions (p < 0.01), but not at 1.00 mg/ml under non-postprandial conditions (Figure 1C).

**Figure 1.**
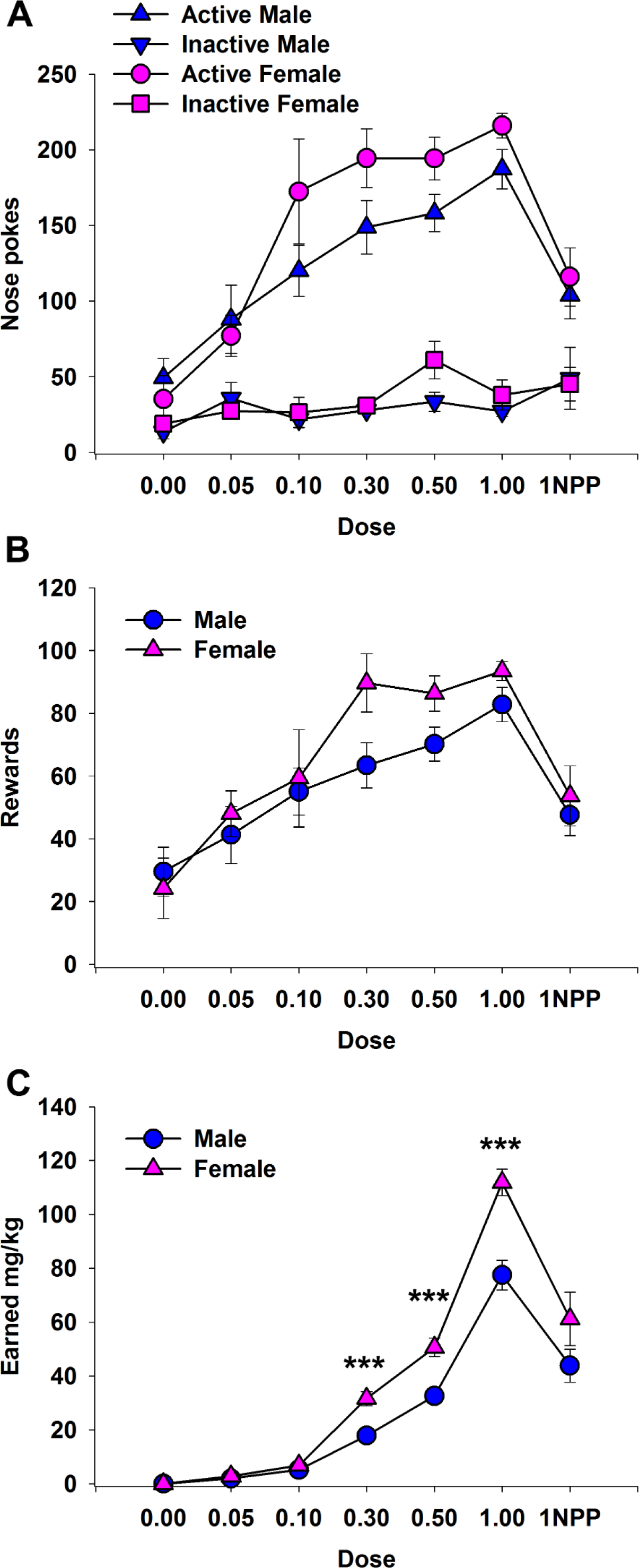
Sex-dependent oral self-administration of oxycodone in mice. Mice orally self-administered different doses of oxycodone over weeks. Active and inactive nosepoke responses (A), rewards earned (B), and earned mg/kg oxycodone (C) were calculated. Female mice showed a dose-dependent significant elevation in earned mg/kg oxycodone compared with male mice. *** p < 0.001 male and female earned mg/kg. Pink – female, blue – male. 1NPP – 1.00 mg/ml dose under non-postprandial conditions. Data are mean ± SEM (error bars).

Following one week of forced abstinence, mice were returned to operant chambers for extinction-reinstatement testing. A between-within cue-induced reinstatement session was used (Shaham et al., 2003) consisting of an extinction phase and a cue phase (Figure 2A). When three or fewer active nosepokes were made during thirty consecutive minutes, the cue phase began. Time to extinction criteria did not differ between sexes, t(14) = 0.69, p > 0.05 (Figure 2B). During the cue phase single active nosepokes activated the tone cue previously associated with drug consumption. Both male and female mice reliably reinstated extinguished oxycodone-seeking behavior following cue presentation (Figure 2C). Active nosepoke rates significantly increased during the cue phase compared with the extinction phase, as reflected by a mixed ANOVA significant main effect of phase, F(1, 14) = 57.23, p < 0.001, with no effect of sex or phase X sex interaction. Inactive nosepoke rates showed no significant main effect of phase, sex, or their interaction.

**Figure 2.**
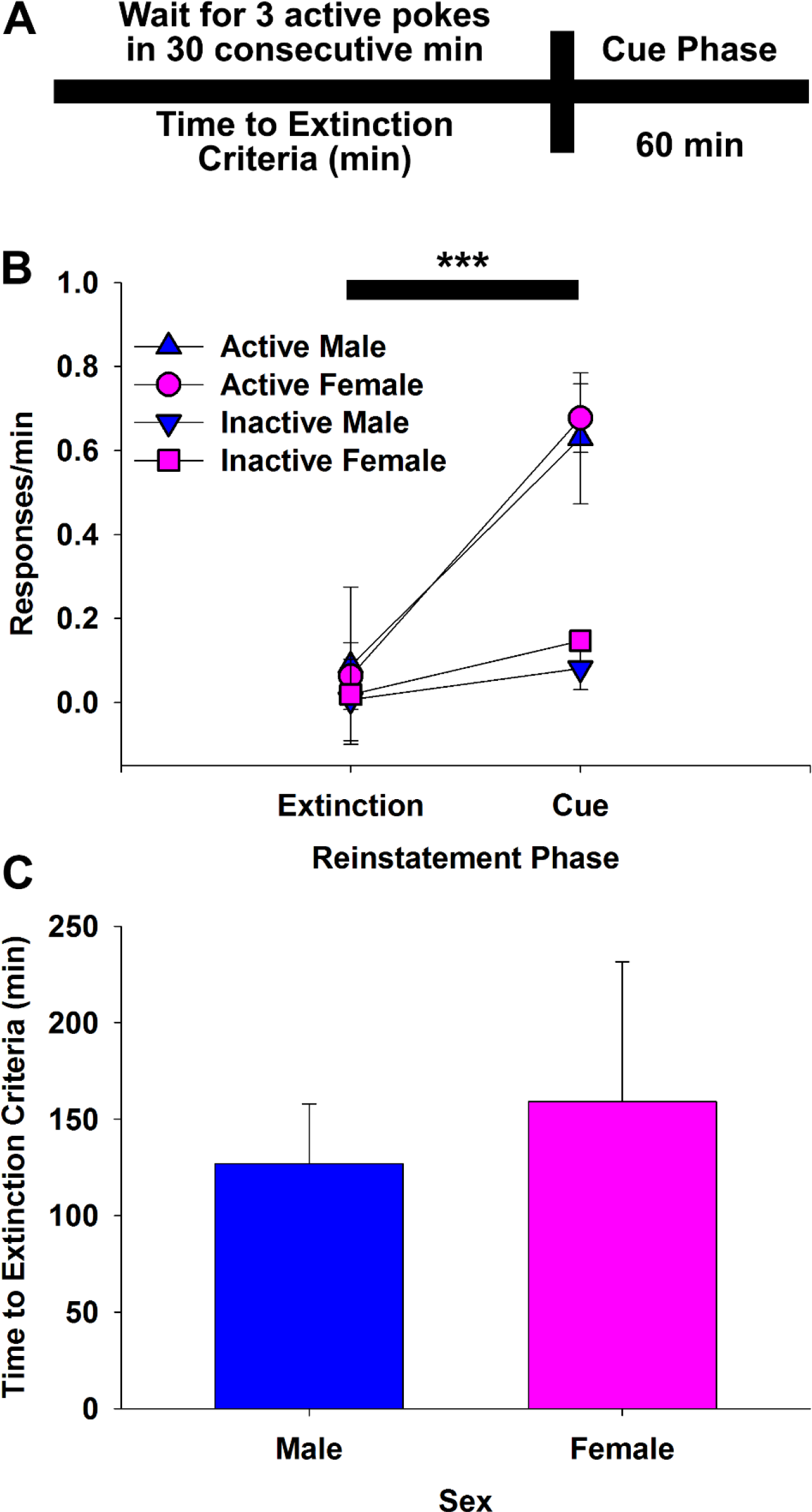
Cue-induced reinstatement of extinguished oral oxycodone-seeking behavior in male and female mice. A. Cue-induced reinstatement timeline. Following one week of abstinence, mice were returned to operant chambers. When mice emitted three or fewer active nosepokes in thirty consecutive minutes, the Time to Extinction Criteria was met, beginning the cue phase. During the cue phase, single active nosepokes activated the cue previously associated with oxycodone reward. B. Male and female mice similarly showed cue-induced reinstatement. C. Male and female mice similarly met time to extinction criteria. *** p < 0.001 between phases for active response rates. Pink – female, blue – male. Data are mean ± SEM (error bars).

Having identified that oral drug cues are sufficient to cause reinstatement of extinguished drug-seeking behavior, we next aimed to identify whether initial water self-administration and postprandial training are required for oral cue-induced reinstatement. A group of male and female mice orally self-administered oxycodone over weeks between 0.05 and 0.10 mg/ml under non-postprandial conditions and did not receive prior water self-administration training. Given that reinstatement did not differ between sexes in our initial experiment (Figure 2C), we pooled male and female mice together. Self-administration was low under non-postprandial conditions. Mice made significantly fewer active nosepokes over doses, z = −2.20, p < 0.05 but did not show a difference in inactive nosepokes, z = −0.10, p > 0.05 (Figure 3A). Consequently, rewards were significantly reduced at 0.10 mg/ml compared with 0.05 mg/ml, z = −2.20, p < 0.05 (Figure 3B). Earned mg/kg oxycodone did not differ between doses, p > 0.05 (Figure 3C). Despite the reduced self-administration of oxycodone under non-postprandial conditions, oral oxycodone cues were sufficient to reinstate extinguished drug-seeking (Figure 3D). Active nosepoke rates were significantly increased during the cue phase compared with the extinction phase, z = −2.20, p < 0.05, while inactive nose poke rates did not differ between phases, p > 0.05. Together, prior water training and postprandial conditions are not required for oral drug cues to acquire incentivizing properties that reinstate responding during abstinence.

**Figure 3.**
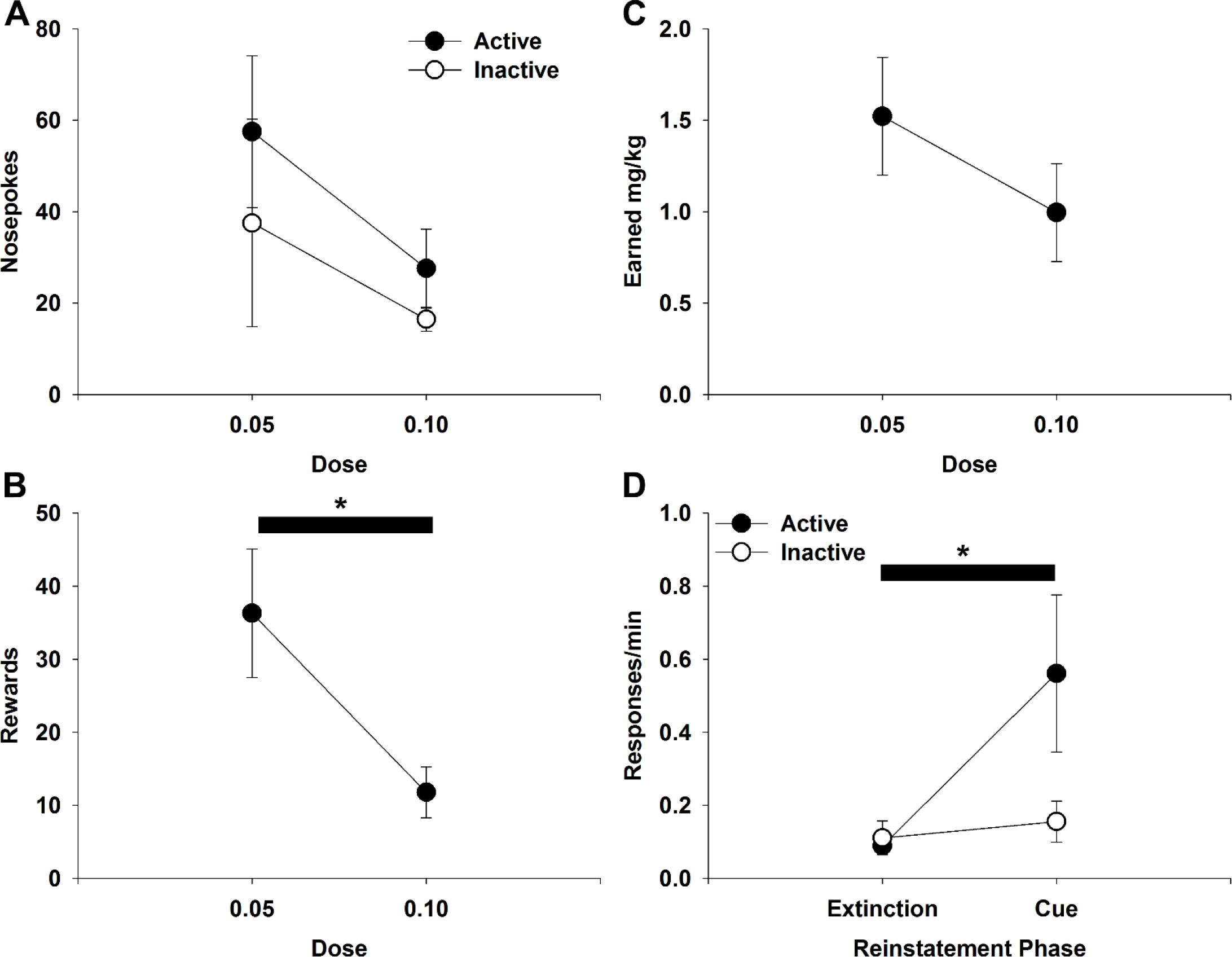
Oral drug-associated cues are sufficient to reinstate extinguished oxycodone-seeking behavior without prior postprandial or water self-administration. Mice orally self-administered different doses of oxycodone. Active and inactive nosepoke responses (A), rewards earned (B), and earned mg/kg oxycodone (C) were calculated. A-C. Mice significantly decreased number of rewards earned across doses. D. Cue-induced reinstatement of extinguished drug-seeking behavior. Active but not inactive nose pokes increased from extinction to cue phases. * p < 0.05. Data are mean ± SEM (error bars).

Given the elevated earned oxycodone in female mice compared with male mice during self-administration training, we examined the effects of oral oxycodone on psychomotor stimulation, drug metabolism, and tested whether intact sex organs influence oral oxycodone self-administration. To investigate differences in psychomotor stimulation and drug metabolism between sexes, mice were orally administered different doses of oxycodone by gavage under non-postprandial and postprandial conditions (Table 2). We first compared psychomotor stimulation under non-postprandial conditions at different doses oxycodone. A 3 × 2 mixed ANOVA yielded a significant dose x sex interaction, F(2, 18) = 27.06, p < 0.001. Posthoc Sidak-adjusted pairwise comparisons showed that both male and female mice had significantly higher speed following 8 or 16 mg/ml oxycodone compared with saline, all p < 0.001. However, female mice had significantly higher speed at saline (p < 0.05), 8 mg/ml oxycodone (p < 0.001), and 16 mg/ml (p < 0.001) (Figure 4A). In addition, females showed a dose-dependent increase in speed from 8 to 16 mg/ml oxycodone (p < 0.001) while male mice did not (p > 0.05) (Figure 4A). Analyzing plasma drug levels, a 2 × 2 mixed ANOVA yielded a significant dose x sex interaction, F(1, 9) = 15.44, p < 0.01. Posthoc Sidak-adjusted pairwise comparisons showed that oxycodone/oxymorphone levels were significantly higher in females compared with males at the 16 mg/ml dose (p < 0.05) but not at the 8 mg/ml dose (p > 0.05) (Figure 4B). In addition, females showed a dose-dependent increase in plasma oxycodone/oxymorphone from 8 to 16 mg/ml oxycodone (p < 0.01) while male mice did not (p > 0.05) (Figure 4B).

**Figure 4.**
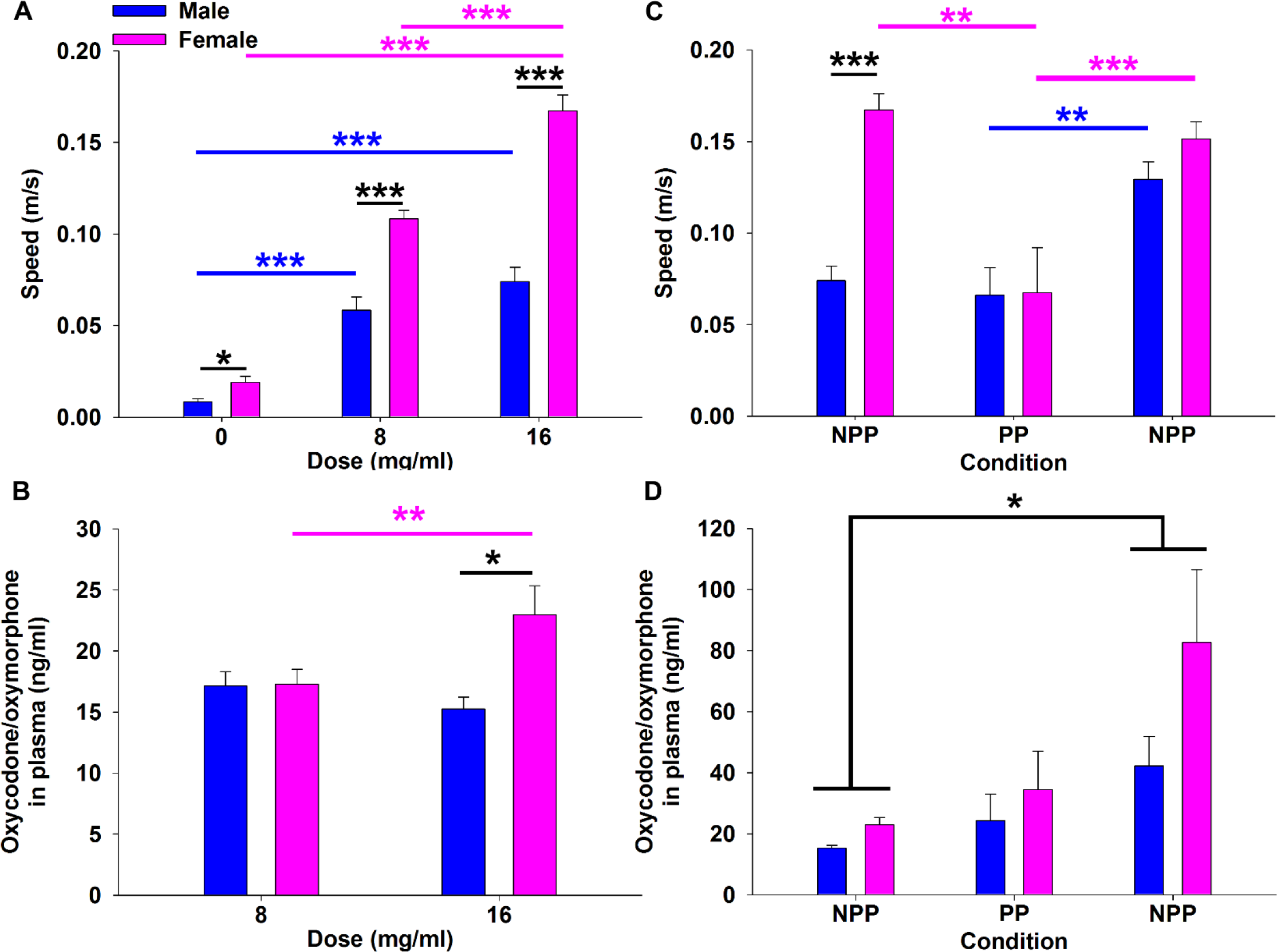
Sex and prandial-dependent psychomotor and metabolic effects of oral oxycodone. A. Mean speed following oral administration of 0 mg/ml, 8 mg/ml, or 16 mg/ml oxycodone. Females were significantly faster across doses. B. Plasma oxycodone/oxymorphone concentration (ng/ml) following oral administration of 8 mg/ml or 16 mg/ml oxycodone. Oxycodone/oxymorphone was significantly greater in females than males selectively following 16 mg/ml administration. C. Mean speed following oral administration of 16 mg/ml oxycodone under non-postprandial (NPP) or postprandial (PP) conditions. Females show greater speed at the initial NPP administration but no differences were observed under PP conditions or the second NPP administration. D. Plasma oxycodone/oxymorphone concentration (ng/ml) following oral administration of 16 mg/ml oxycodone under NPP or PP conditions. Oxycodone/oxymorphone significantly increased between the first and second non-postprandial administration in both sexes. Blue asterisks indicate within male comparison. Black asterisks indicate between sex comparison. *** p < 0.001. ** p < 0.01. * p < 0.05. Pink – female, blue – male. Data are mean ± SEM (error bars).

We continued by comparing psychomotor stimulation and plasma oxycodone/oxymorphone under different prandial conditions at 16 mg/ml oxycodone in an ABA design (Table 2). Locomotion analyzed in a 3 × 2 mixed ANOVA yielded a significant condition X sex interaction, F(2,18) = 9.279, p < 0.01. Posthoc Sidak-adjusted pairwise comparisons showed that prandial administration significantly reduced speed for female mice (p < 0.01) but not male mice (Figure 4C). However, after returning to non-postprandial administration both male and female mice showed significantly higher speed than under postprandial conditions (all p < 0.01) (Figure 4C). Whereas female mice showed significantly higher speed at the first non-postprandial oral dose (p < 0.001), speed did not differ between sexes at the second non-postprandial oral dose (p > 0.05) (Figure 4C). Plasma drug levels analyzed in a 3 x 2 mixed ANOVA yielded a significant main effect of condition, F(2, 18) = 8.884, p < 0.01, with no significant main effect of sex or sex X condition interaction. Posthoc Sidak-adjusted pairwise comparisons showed that oxycodone/oxymorphone levels were significantly higher following the second non-postprandial administration compared with the first (p < 0.05) (Figure 4D). Together, these results suggest that female mice are more sensitive to the psychomotor stimulation aspects of oral oxycodone than male mice. However, with extended oxycodone experience psychomotor stimulation does not differ between sexes. Interestingly, the psychomotor response of mice following oral oxycodone was not always predicted by changes in plasma oxycodone/oxymorphone. Further, the increase in plasma oxycodone/oxymorphone in both sexes by repeated oxycodone exposure suggests that drug metabolism is altered by repeated exposure.

To test the hypothesis that the difference in earned drug between male and female mice was the result of intact sex hormone-producing testes and ovaries, male and female mice received either gonadectomy or sham surgical procedures. Ovariectomized and sham mice orally self-administered escalating doses of oxycodone over weeks (Table 3). For female mice, a mixed ANOVA yielded a significant interaction of surgery X dose for active nosepokes, F(6, 96) = 3.44, p < 0.01. Sidak-adjusted pairwise comparisons indicated that ovariectomized mice made significantly more active nosepokes than sham mice selectively under non-postprandial conditions, p < 0.05 (Figure 5A). No main effect of surgery, dose, or their interaction was yielded for inactive nosepokes (Figure 5A). Similar to active nosepokes, ovariectomized mice earned significantly more rewards than sham mice selectively under non-postprandial conditions, surgery x dose interaction: F(6, 96) = 2.83, p < 0.05; Sidak-adjusted pairwise comparison: p < 0.05 (Figure 5B). While a significant main effect of dose was yielded for earned mg/kg oxycodone, F(6, 96) = 23.471, p < 0.001, there was no significant main effect of surgery or interaction of dose and surgery (Figure 5C). Following non-postprandial self-administration, mice self-administered oral oxycodone at 1.0 mg/ml under non-postprandial conditions with a progressive ratio 4 schedule of reinforcement. Breakpoints did not significantly differ between ovariectomized and sham groups, Mann-Whitney Test, z = −1.63, p > 0.05 (Figure 5D).

**Figure 5.**
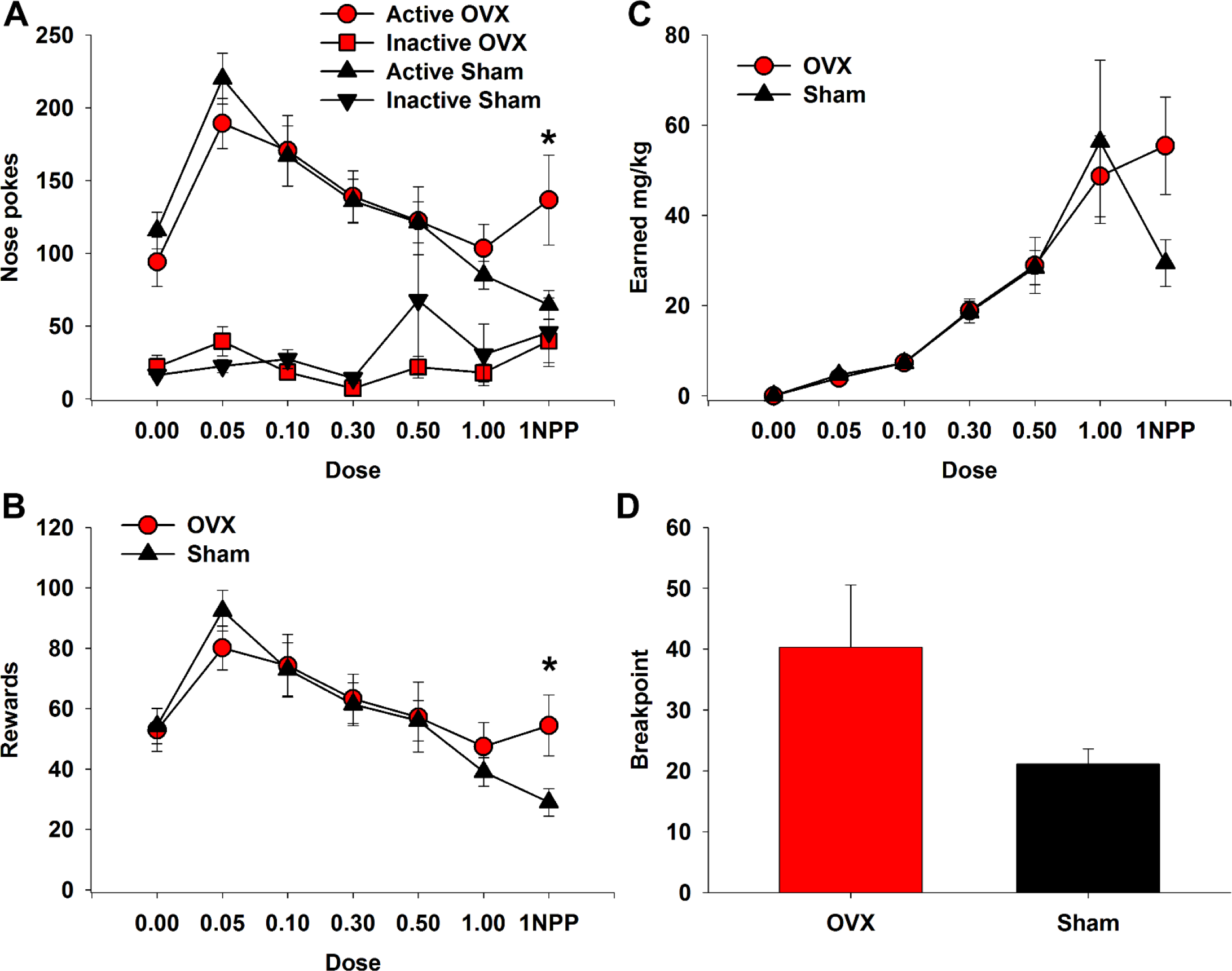
Ovariectomy dose-dependently enhances oral oxycodone self-administration. Mice orally self-administered different doses of oxycodone over weeks. Active and inactive nosepoke responses (A), rewards earned (B), and earned mg/kg oxycodone (C) were calculated. A-C. Ovariectomy mice showed a dose-dependent significant elevation over sham mice in active nose pokes and rewards selectively during non-postprandial self-administration of 1.00 mg/ml oxycodone. D. No significant difference in breakpoints between ovariectomy and sham mice. * p < 0.05. 1NPP – 1.00 mg/ml dose under non-postprandial conditions. OVX – ovariectomy. Red – ovariectomy, black – sham. Data are mean ± SEM (error bars).

We next examined orchiectomized and sham male mice for differences in oral oxycodone self-administration. Prior to beginning non-postprandial self-administration, one sham mouse died of overdose. To statistically analyze differences between orchiectomy and sham male groups with a missing datapoint in one sham mouse’s non-postprandial data, we utilized linear mixed models that do not require subjects to have the same number of observations. Examining active nosepokes, the linear mixed model yielded significant main effects of dose, F(6, 71) = 18.10, p < 0.001, and surgery, F(1, 12) = 5.09, p < 0.05, but no significant dose X surgery interaction. This result indicates that sham mice made significantly more active nosepokes than orchiectomy mice regardless of dose (Figure 6A). No main effect of surgery, dose, or their interaction was yielded for inactive nosepokes (Figure 6A). Sham mice earned significantly more rewards than orchiectomy mice at all doses, main effects of dose, F(6, 71) = 26.70, p < 0.001, and surgery, F(1, 12) = 7.07, p < 0.05, but no significant surgery X dose interaction (Figure 6B). While a significant main effect of dose was yielded for earned mg/kg oxycodone, F(6, 71) = 21.54, p < 0.001, there was no significant main effect of surgery or interaction of dose and surgery (Figure 6C). There was also no significant difference in breakpoints under a progressive ratio 4 schedule of reinforcement, t(11) = −0.55, p > 0.05 (Figure 6D).

**Figure 6.**
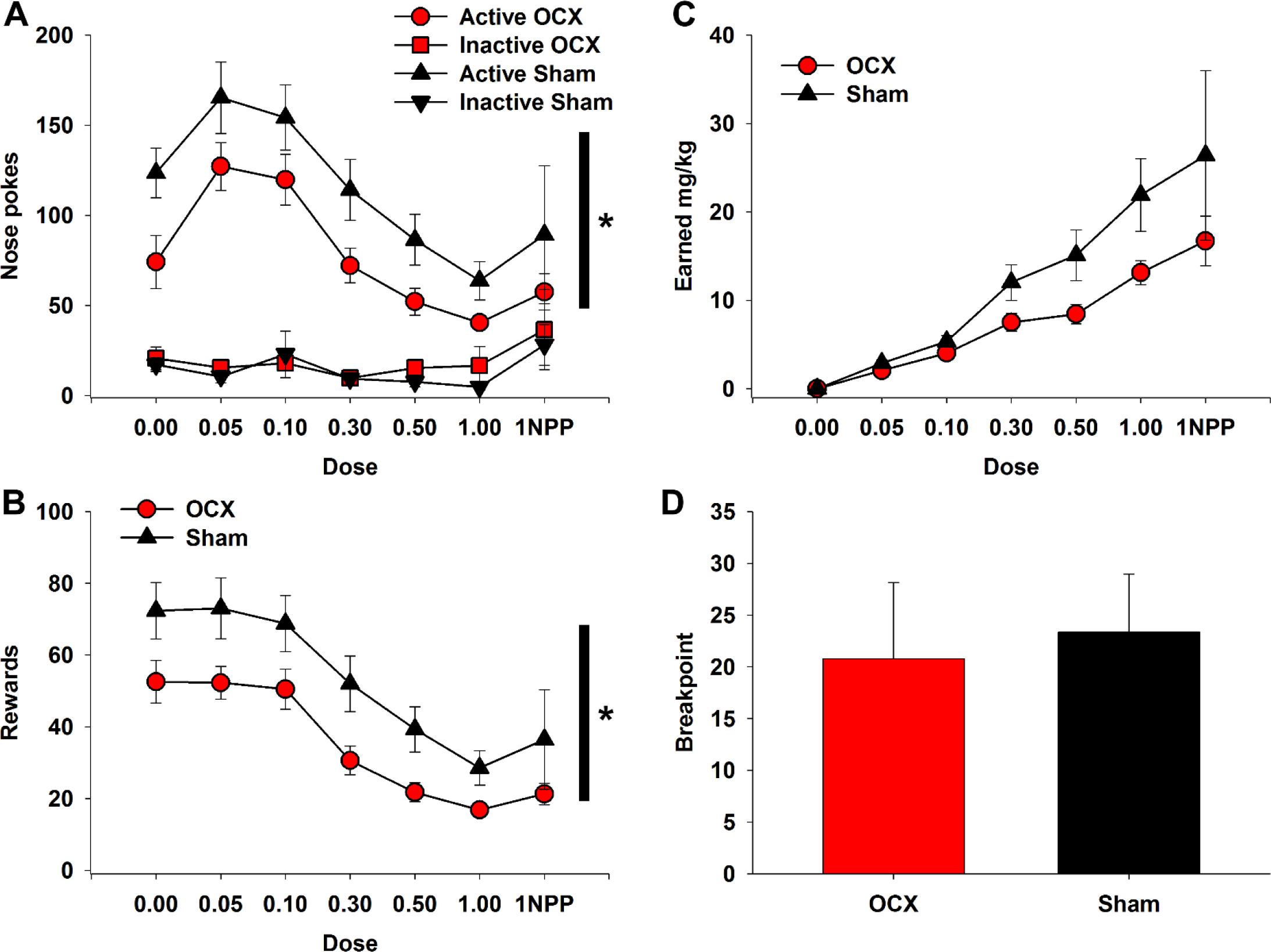
Orchiectomy blunts oral oxycodone self-administration. Mice orally self-administered different doses of oxycodone over weeks. Active and inactive nosepoke responses (A), rewards earned (B), and earned mg/kg oxycodone (C) were calculated. A-C. Orchiectomy mice showed a significantly decreased number of active nose pokes and rewards earned across doses. D. No significant difference in breakpoints between orchiectomy and sham mice. * p < 0.05. 1NPP – 1.00 mg/ml dose under non-postprandial conditions. OCX – orchiectomy. Red – orchiectomy, black – sham. Data are mean ± SEM (error bars).

Comparing the male and female sham groups, we replicated that female mice orally self-administered more than male mice. A linear mixed model yielded a significant dose X surgery interaction for active responses, F(6, 83) = 3.14, p < 0.01. Posthoc Sidak-adjusted pairwise comparisons showed that female mice made significantly more active nosepokes at 0.05 mg/ml but not under other doses or conditions, p < 0.05 (Figure 7A). No main effect of surgery, dose, or their interaction was yielded for inactive nosepokes (Figure 7A). While a significant dose X surgery interaction was yielded from number of rewards, F(1, 83) = 3.72, p < 0.01, posthoc Sidak-adjusted pairwise comparisons did not identify differences between groups that reached statistical significance at any dose (Figure 7B). However, a significant surgery X dose interaction was yielded for earned mg/kg oxycodone, F(6, 83) = 3.27, p < 0.01. Sidak-adjusted pairwise comparisons showed that female mice had significantly higher earned mg/kg oxycodone between 0.50 and 1.00 mg/ml doses under postprandial conditions, but not at 1.00 mg/ml under non-postprandial conditions, all p < 0.01 (Figure 7C). There was no significant difference in breakpoints under a progressive ratio 4 schedule of reinforcement, t(13) = −0.39, p > 0.05 (Figure 7D).

**Figure 7.**
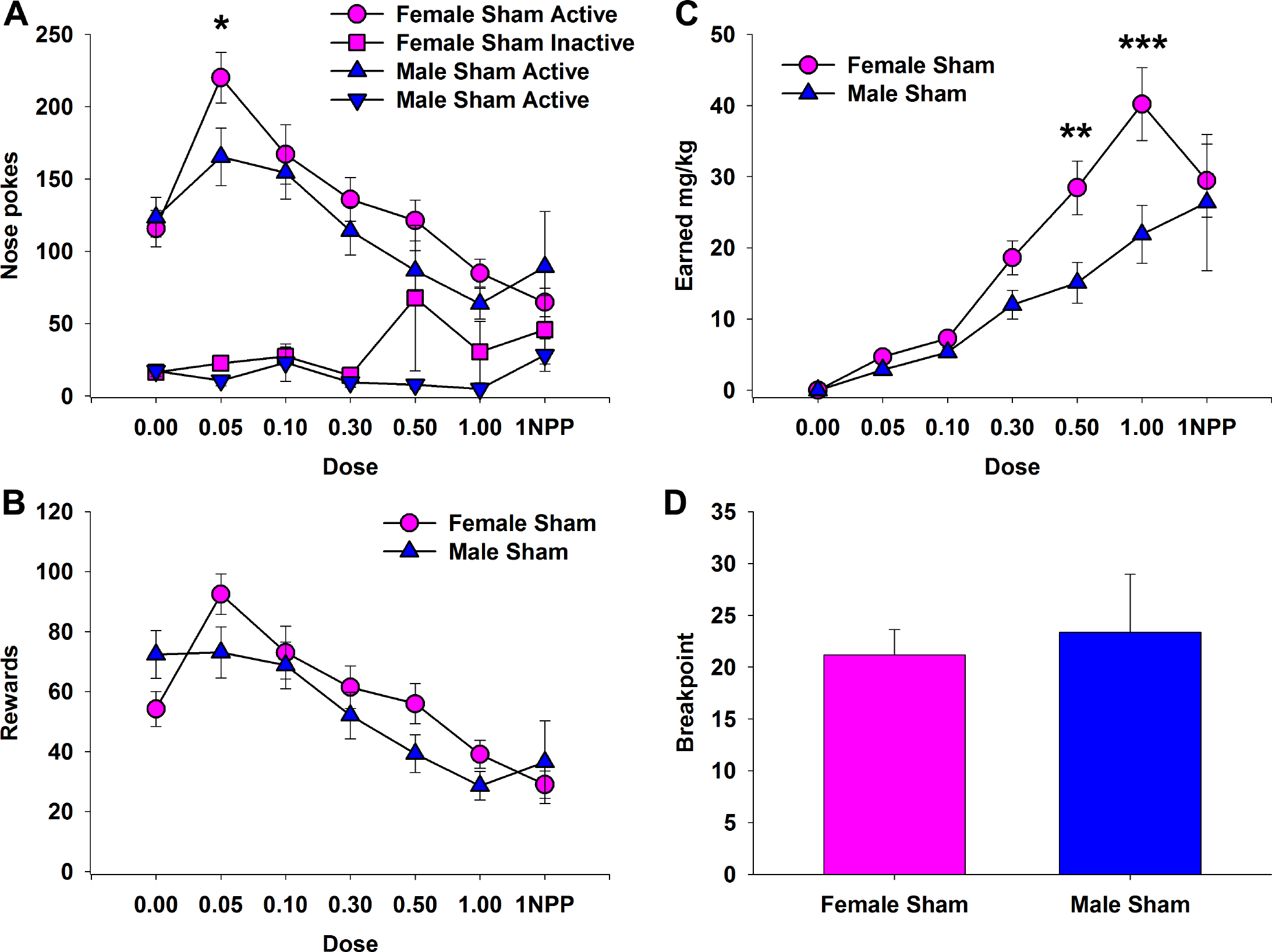
Sex-dependent oral self-administration of oxycodone in sham mice. Mice orally self-administered different doses of oxycodone over weeks. Active and inactive nosepoke responses (A), rewards earned (B), and earned mg/kg oxycodone (C) were calculated. A-C. Female sham mice showed a dose-dependent significant elevation over male sham mice in active nose pokes at the lowest concentration and significantly elevated earned mg/kg oxycodone at 0.50 and 1.00 mg/ml oxycodone concentrations. D. No significant difference in breakpoints between male and female sham mice. *** p < 0.001. ** p < 0.01. * p < 0.05. 1NPP – 1.00 mg/ml dose under non-postprandial conditions. Pink – female sham mice, bleu – male sham mice. Data are mean ± SEM (error bars).

## Discussion

Oral consumption is an increasingly popular route of administration by prescription opioid users (Back, Lawson, Singleton et al., 2011; Gasior, Bond & Malamut, 2016; Kirsh, Peppin & Coleman, 2012; Surratt, Kurtz & Cicero, 2011; Young, Havens & Leukefeld, 2010). However, animal models of opioid addiction have focused on intravenous routes, limiting understanding of a significant population of opioid users that consume orally. Here, we used a modified version of the recently established oral model of oxycodone self-administration in mice (Enga, Jackson, Damaj et al., 2016) to evaluate whether oral drug cues are sufficient to cause relapse, and whether sex differences exist in the susceptibility to relapse, as modeled in the extinction-reinstatement paradigm.

We found that oral drug cues reliably reinstated extinguished drug-seeking behavior in both male and female mice. These data support the observation that both oral and intravenous-associated drug cues are capable of causing cravings in human users (McHugh, Fulciniti, Mashhoon et al., 2016). In addition, these data indicate that drugs associated with oral ingestion, despite having less immediate effects than intravenous use (Gasior, Bond & Malamut, 2016; Kirsh, Peppin & Coleman, 2012; Leow, Smith, Watt et al., 1992), are nonetheless capable of invigorating drug-seeking behavior. A potentially problematic aspect of the oral model of oxycodone self-administration in mice is that the initial learning of self-administration behavior uses water reward and that most of self-administration training takes place under postprandial conditions. However, we found that without prior water self-administration training and without postprandial conditions, oral drug cues were still sufficient to reinstate extinguished drug-seeking behavior. Therefore, prior water and prandial training are not required for oral drug cues to cause reinstated drug-seeking behavior.

Mixed results have been reported in the literature regarding the presence and direction of sex differences in opioid-related behavior (Collins, Reed, Zhang et al., 2016; Lynch, Roth & Carroll, 2002; Mavrikaki, Pravetoni, Page et al., 2017; Serdarevic, Striley & Cottler, 2017). We did not observe a sex difference in reinstatement. However, we found that in both our initial examination of oxycodone self-administration and comparing sham male and sham female mice in subsequent experiments, that female mice earned significantly more mg/kg oral oxycodone than male mice. These results support clinical observations that female opioid users are more likely to advance to regular prescription opioid use than males (Back, Lawson, Singleton et al., 2011; Gasior, Bond & Malamut, 2016; Kirsh, Peppin & Coleman, 2012). Additionally, these findings support results in rats showing females have higher intravenous oxycodone intake compared with males (Mavrikaki, Pravetoni, Page et al., 2017). The oral oxycodone model may therefore be an additional avenue for modeling female prescription opioid users.

To identify potential factors that influenced the larger earned mg/kg oxycodone in females compared with males, we examined psychomotor stimulation and drug metabolism from plasma concentrations of oxycodone/oxymorphone following experimenter-administered oral oxycodone. Subsequently, we examined the role of intact gonads on oral self-administration. After initial exposure to 8 mg/ml and 16 mg/ml oral oxycodone, female mice showed significantly higher psychomotor stimulation than male mice. The higher psychomotor stimulation of female mice at the lower oxycodone dose might explain the significantly higher responding of female sham mice at the lowest oxycodone dose compared with male sham mice (0.05 mg/ml). That is, enhanced responding at this dose by female mice may reflect a more reinforcing effect than males. Interestingly, despite the enhanced psychomotor stimulation of females at the lowest oral dose, plasma oxycodone/oxymorphone concentrations did not differ between sexes. We nonetheless found that at the first 16 mg/ml experimenter-administered oral oxycodone dose females showed both significantly higher psychostimulation and plasma oxycodone/oxymorphone compared with males. Further, while females showed dose-dependent increases in both psychostimulation and plasma oxycodone/oxymorphone concentrations, males did not. Together, these results suggest that oral use has larger psychostimulation and metabolic effects in females than males, which may support their larger use in human female users.

Prandial conditions had complex effects on psychostimulation and oxycodone/oxymorphone plasma concentrations. While females showed significantly decreased psychomotor stimulation following postprandial oral administration, males did not. Further, both male and female oxycodone/oxymorphone plasma concentrations did not differ between non-postprandial and postprandial conditions. Instead, plasma oxycodone/oxymorphone concentrations increased in both males and females across each subsequent administration, whether postprandial or not. We had attempted to design our experiment to limit the development of tolerance, which can rapidly form after four days of twice-daily oxycodone oral administrations (Jacob, Poklis, Akbarali et al., 2017), by administering one dose at least three days apart from the next. Nevertheless, the significant increase in plasma oxycodone/oxymorphone between the first and second non-postprandial 16 mg/ml oral dose suggests that metabolic changes have occurred between these doses. In addition, at the second non-postprandial oral dose of 16 mg/ml, males showed equal psychomotor stimulation as females, whereas male psychomotor stimulation was significantly less than females following the first non-postprandial 16 mg/ml oral dose. We interpret these results in the following manner. First, postprandial oxycodone delivery diminishes female psychostimulation more readily than males, but with repeated exposure male postprandial psychostimulation is significantly decreased compared to non-postprandial administration. Second, oxycodone/oxymorphone metabolism does not always predict changes in psychostimulation, suggesting that other factors, for instance the binding of oxycodone/oxymorphone to central μ opioid receptors, may be altered by repeated exposure to oral oxycodone.

As predicted by the results of the experimenter-administered oral oxycodone experiments, our results suggest that the role of sex is complex in oral oxycodone self-administration. While female gonadectomized mice did not differ from female shams in self-administration under postprandial conditions, female gonadectomized mice responded significantly more than sham mice at the highest dose under non-postprandial conditions. The higher psychomotor stimulation of females under non-postprandial conditions compared with postprandial conditions may explain why sham female mice decreased self-administration between 1.00 mg/ml postprandial and 1.00 mg/ml non-postprandial doses. The increased responding of ovariectomy mice at 1.00 mg/ml non-postprandially to 1.00 mg/ml postprandially, which differed from shams, suggests that ovariectomy may have reduced the rewarding aspects of non-postprandial oxycodone at the 1.00 mg/ml dose. Together with the subsequent observation that ovariectomized mice did not differ from shams under a progressive ratio schedule of reinforcement under the same dose and non-postprandial conditions suggests that a complex interaction of sex hormones, effort, and reward motivation, likely influenced by psychostimulation, regulates oral self-administration of oxycodone in female mice.

In contrast to females, the effects of gonadectomy on oral oxycodone self-administration were more dramatic in males. Oral oxycodone self-administration was blunted in male gonadectomized mice across all doses and prandial conditions compared to sham males. Nevertheless, we again did not observe a significant difference between gonadectomized and sham male mice under a progressive ratio schedule of reinforcement. Together, these results suggest that prandial conditions may play a role in oral oxycodone self-administration and that the prandial influence on oxycodone intake may be more susceptible in females than males. This suggestion is supported by 1) the significantly reduced psychomotor stimulation of female mice, but not male mice, following initial exposure to oral oxycodone and 2) both orchiectomized and sham male mice maintained or increased responding at 1.00 mg/ml non-postprandially compared with 1.00 mg/ml postprandially. It is noted though that in intravenous self-administering rats, food restriction strongly influences opioid-seeking behavior (Shalev, 2012). Whether similar mechanisms occur between the role of food restriction on intravenous opioid-seeking and the role of postprandial thirst on oral opioid-seeking remains to be determined.

It is important to consider that our study identified unexpected variability between different cohorts of oral self-administering mice. The dose-response and dose-reward curves of our initial study were different from all subsequent examinations. In our initial study both male and female mice increased or maintained high rates of self-administration as the dose increased. This was unexpected and in contrast to the results reported in the initial oral oxycodone self-administration paradigm (Enga, Jackson, Damaj et al., 2016). All follow up experiments showed a decreasing dose-response and dose-reward relationship, including under non-postprandial conditions. The difference between our initial and subsequent examinations of oral oxycodone self-administration are unclear. However, we did find in our initial cohort a significant amount of leftover oxycodone in the magazine that was uncommon for subsequent examinations. This may account for the higher earned mg/kg oxycodone in our initial examination. The potential for unconsumed oxycodone is an important difference between oral and intravenous paradigms of oxycodone self-administration. Therefore, the oral oxycodone paradigm may offer a unique opportunity to determine the factors that delineate responding for and consuming earned oxycodone. Finally, without constraints of jugular catheterization patency, the oral oxycodone self-administration paradigm has desired qualities for experimentation using protracted self-administration thought important for addiction. A major benefit of the mouse model is the large array of genetic tools for manipulation. The present results provide a critical first step toward these experiments by providing a platform to identify the behavioral, genetic, and molecular bases of cued relapse involved in oral opioid-seeking behavior.

## Acknowledgements

This research was supported by the Boettcher Foundation’s Webb-Waring Biomedical Research Awards program and The University of Colorado. We thank Dr. Serge Campeau for technical assistance with ELISA and Drs. Zoe Donaldson and Deena Walker for advice on gonadectomy procedures. The funders had no role in study design, data collection and analysis, decision to publish, or preparation of the manuscript. The authors have no financial interests to be disclosed.

## Author contributions

AGP, DJM, and DHR were responsible for the study concept and design. AGP, DJM, SL, KR, DTH, SKE, KNF, and DHR contributed to the acquisition of animal data. AGP, DJM, SL, and DHR performed data analysis. AGP, DJM, and DHR wrote the manuscript with the contribution of all authors.

